# Autonomous combinatorial color barcoding for multiplexing single molecule RNA visualization

**DOI:** 10.1101/127373

**Authors:** Yong-Sheng Cheng, Yue Zhuo, Katharina Hartmann, Peng Zou, Gözde Bekki, Heike Alter, Hai-Kun Liu

## Abstract

Single molecular fluorescence in situ hybridization (smFISH) detects RNA transcripts with spatial information and digital molecular counting. However, the broad usage of smFISH is still hindered by the complex chemical probe conjugation or microscopy set-up, especially for investigating multiple gene expression. Here we present a multiple fluorophore enzymatic labeling method (termed HuluFISH) for smFISH probes to achieve flexible combinatorial color barcoding in single hybridization step. The multiplex capacity of HuluFISH follows an exponential growth with the increase of the number of fluorophore types. We demonstrate that this method can be used to detect cellular heterogeneity in embryonic mouse brain on single cell level.

## Introduction

Since the invention of *in situ* hybridization^1^,it has been continuously advancing our understanding of gene expression with spatial information. The smFISH technology pioneered by Robert Singer’s lab^2^ and further developed by Raj et al.^3^, brings *in situ* RNA quantification into single molecular and digital manner. Nevertheless, the limited choices of single fluorophore on probes cannot cope with the increasing demand of simultaneous multiple gene detections. Although sequential hybridization has been employed to achieve multiplex gene detection using smFISH^4,5^, sophisticated experimental settings hinder its broad applications in the biomedical community. One alternative strategy for increasing the multiplexity beyond fluorophore limit is using combinatorial color barcoding via spectral or spatial separated groups of smFISH probe^6,7^. Current combinatorial barcoding either needs a long gene target for mRNA^6^ or only targets intrinsically non-stable introns^7^, and thus restricts their applications from detecting the majority of transcripts (median mouse coding DNA sequence (CDS) length is 1026 bp). Therefore, a combinatorial color barcoding on the individual smFISH probe will empower the conventional smFISH with massive color combinations with the same number of probes.

## Results

Conventional smFISH probes or its derivatives are using chemical labeling for conjugating a fluorophore to the internal, 5’ or 3’ end of an unlabeled oligonucleotide pre-equipped with an amine group, which is readily reacting with fluorophores functionalized with a N-Hydroxysuccinimide (NHS) ester^2,8,9^. We apply a novel enzymatic fluorophore labeling method, which is based on the usage of T4 DNA ligase (T4DL), for HuluFISH 1.0. It does not require any amine modification of unlabeled gene-specific oligonucleotides (GSO) for HuluFISH probe. As a consequence, it is now possible to cost-effectively synthesize single fluorophore labeled smFISH probes (Figure 1a). Comparing with other enzymatic labeling methods we have tested, T4DL has the most cost-effective design (Suppl. Figure1a). In the T4DL based labeling strategy, only a standard PCR primer quality oligonucleotide is required, and the free 3’ hydroxyl group from the GSO is enzymatically conjugated with a common pre-fluorescently-labeled oligonucleotide (termed Hulu), mediated by an adaptor with 4 bp 3’ degenerative sequence to facilitate duplex formation (Figure 1a). This new T4DL based chemistry also abolishes the necessity of HPLC purification of the HuluFISH probe (Suppl. Figure1b and 1c). The polyacrylamide gel electrophoresis (PAGE) purified mouse Gapdh probe has comparable detection sensitivity with the commercially available one (Figure 1b).

**Figure 1:**
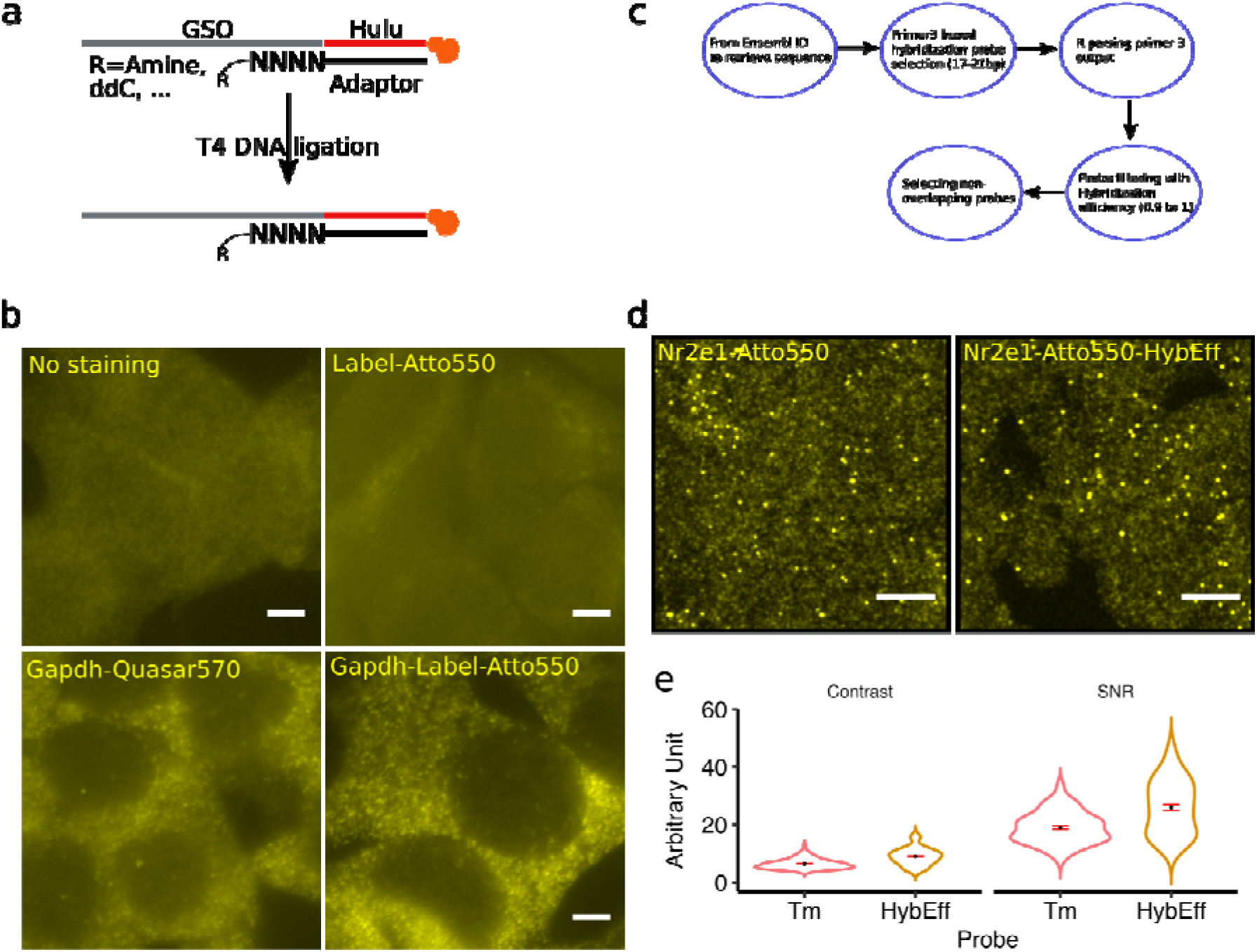
HuluFISH 1.0 probe’s enzymatic labeling and improved probe design. (**a**) T4DL based labeling scheme for HuluFISH probe with standard 3’ end -OH group of GSO. (**b**) *in situ* staining with HuluFISH probe (Gapdh-Label-Atto550) and commercial mouse Gapdh probe (Gapdh_Qusar570) in Hepa 1-6 cell. (**c**) smFISH probe selection pipeline used for all following probes in this paper. Scale bar: 10 *µ*m. (**d**) smFISH detection of low-expressing gene, Nr2e1 in embryonic brain tissue with conventional Tm based or our new hybridization efficiency based probe design. Scale bar: 5 *µ*m. (**e**) Contrast and SNR analysis for Nr2e1’s conventional (Tm) and our (HybEff) design. Between Tm and HybEff, Wilcoxon test’s p-value for contrast and SNR are 8.9 × 10^-10^ and 8.5 × 10^-7^.(n=92, 83 respectively)

Currently, the smFISH probe selection is based on melting temperature (Tm)^3^ or Gibbs free energy^10^, which are not very indicative of probe hybridization efficiency. We developed a pipeline based on Primer3^11^ and DECIPHER^12^ to design and filter for GSO with high hybridization efficiency, which is a more tangible indicator (Figure 1c). Comparing with the conventional Tm based method, our probe design has a better signal-to-noise ratio (SNR) and higher contrast (Figure 1d and 1e). With this new approach, we still have short probe (17-21 bp) to minimize the off-target effect, and a good balance between hybridization capacity and the number of probes we could design for smaller RNA (minimally 24 GSOs). This can be used for customized probe design for any other smFISH methods.

In principle, our T4DL based labeling method also enables multiple fluorophore labeling if the Hulu oligonucleotide is pre-synthesized with multiple fluorophores. However, the technical complexity increases with the number of fluorophores to be incorporated into a single oligonucleotide. Therefore, we extended the T4DL based labeling to multiple-way ligation for incorporating multiple single fluorophore labeled Hulu oligonucleotides (HuluFISH 2.0, Figure 2a). Ligation control experiment shows that HuluFISH 2.0 has a specific ligation product for Gapdh HuluFISH probes, and higher yield compared with the HuluFISH 1.0 (Suppl. Figure 2a). The HuluFISH 2.0 is insensitive to ligation conditions (Suppl. Figure 2b), which demonstrates the robustness of its probe preparation over temperature, reaction time, etc. One critical challenge for using multi-colored probe is that when multiple fluorophores are close to each other, they could be quenched by multiple mechanisms, for example, self-quenching and Förster resonance energy transfer (FRET)^13,14^. Considering the size limitation of the Hulu oligonucleotide, here we use 15 bp spacing for the individual dye, and an adaptor oligonucleotide annealed with the Hulu oligonucleotide in order to rigidify the ssDNA backbone for dyes (Figure 2b).

**Figure 2:**
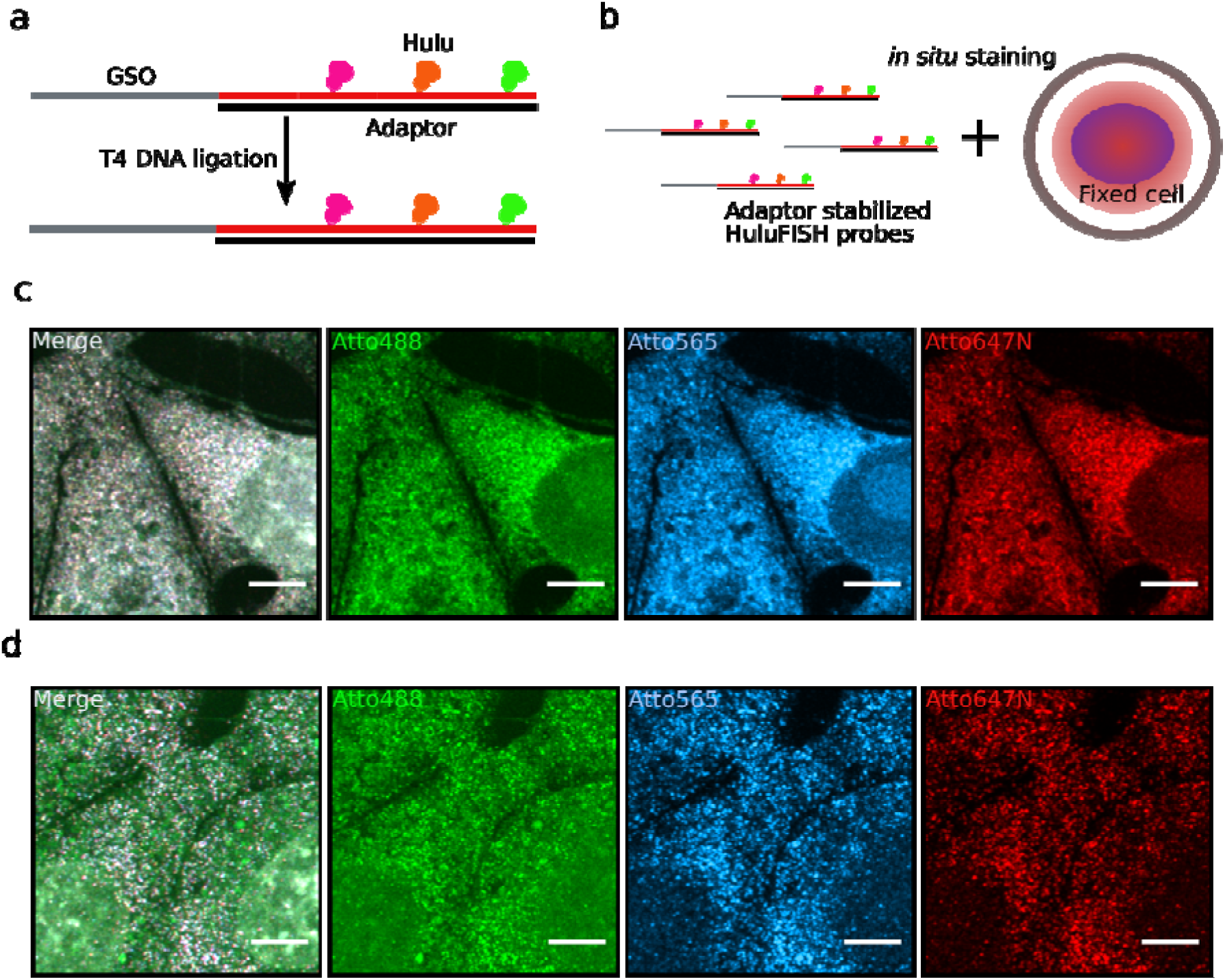
Multiple fluorophore labeling based on HuluFISH 2.0. (**a**) Multiple-way ligation based fluorophore labeling of HuluFISH probes. (**b**) *in situ* staining with HuluFISH probes pre-annealed with the adaptor to avoid multiple fluorophore quenching. (**c**) Gapdh expression in Hepa 1-6 with the Gapdh HuluFISH probe conjugated with Atto488, Atto565, and Atto647N, without adaptor stabilization. (**d**) Gapdh mRNA visualized as individual dots by adaptor pre-annealed HuluFISH probe. Scale bar is 5 *µ*m.

Gapdh probe staining without the adaptor masks the FISH signal by fusing dots with a high background in all channels for Atto488, Atto565, and Atto647N (Figure 2c). And these dot-like signals in 3 channels are not co-localized very well. With the stabilization by the adaptor oligonucleotide, individual clear dots can be obtained in all 3 channels and well co-localized within every channel for Gapdh probes (Figure 2d). Without GSO, the Hulu-adaptor duplex does not generate any dot-like signal (Suppl. Figure 2c). With the multiple labeling capacity of our method, we could assign various color combinations to a panel of genes, and decode the dots by counting their appearance in channels (Suppl. Figure 2d). The evolved multiple fluorophore labeling capability with HuluFISH 2.0 extends the conventional smFISH with an autonomous combinatorial color barcoding mechanism. Fluorophores in each color combination are covalently linked with individual probe, therefore the fluorophore stoichiometry is invariable between probes. During imaging acquisition, the intensity ratio between fluorophores will be independent of the brightness of FISH dots. The barcoding capacity simply increases with the exponentials of the channel (fluorophore choice) number n (the theoretical number of combinations is the sum of all color combinations: 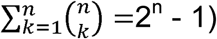. If the relative ratio of the maximal intensities of each FISH dot among channels can be precisely determined, the number of combinations can be higher.

One of the most interesting applications for smFISH is exploring the multiple gene expression patterns in tissue samples. Just with 3 base colors, the color combinations can be used to detect 7 genes in one round of hybridization. Here we use embryonic day 12.5 (E12.5) mouse telencephalon cryo-section samples to visualize the tissue heterogeneity of these 7 genes (Figure 3a). Simultaneous 7-gene detection shows the molecular heterogeneity of fetal brain neural progenitors *in vivo* (Figure 3b). Hierarchical cluster analysis reveals subgroups of mouse telencephalon neural progenitors on single cell resolution (Figure 3c and 3d).

**Figure 3:**
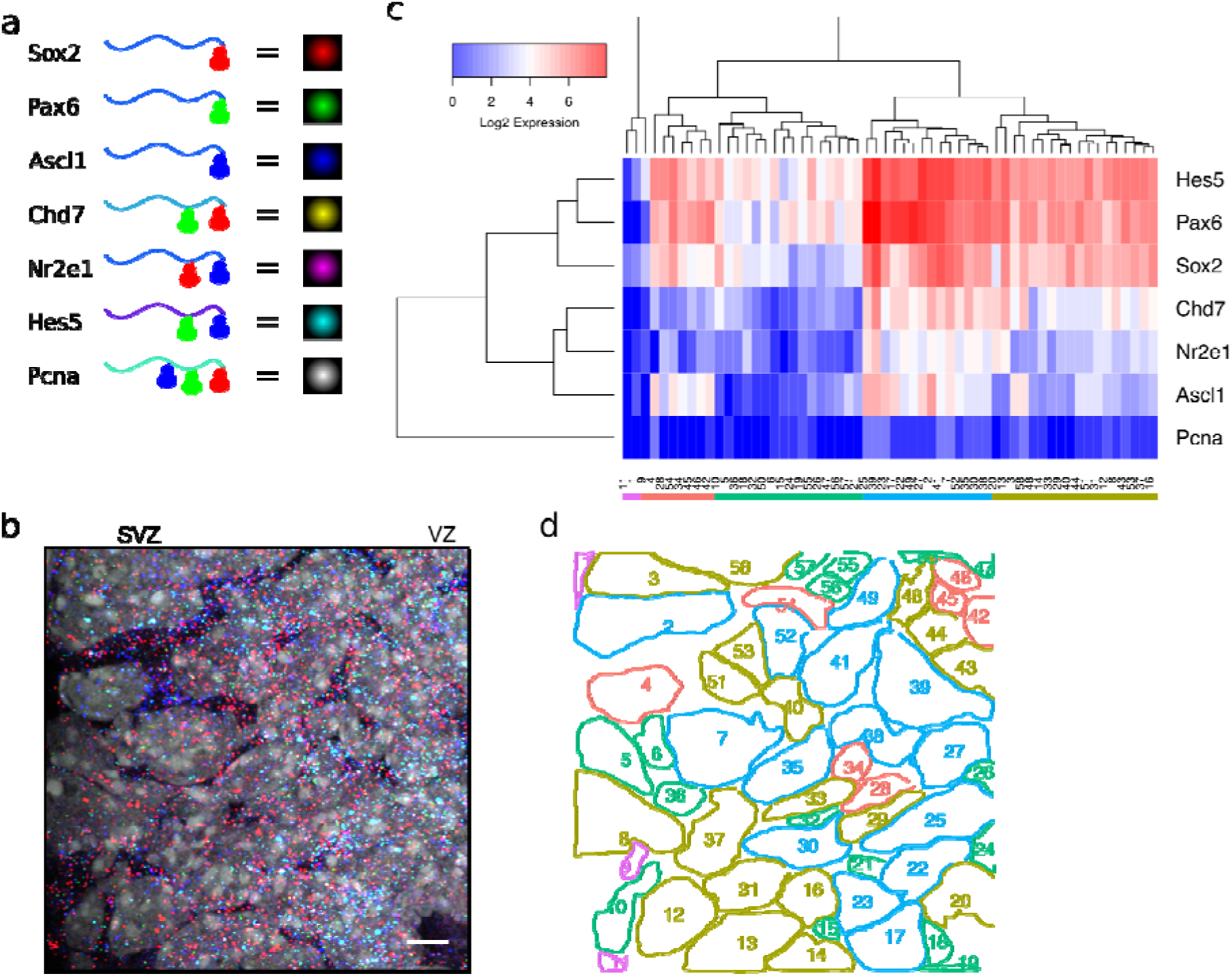
HuluFISH detection of 7 genes in mouse embryonic brain. (**a**) Color coding scheme for HuluFISH from 3 base colors. (**b**) 7-gene detection in E12.5 mouse embryonic brain ventricular zone (VZ) and subventricular zone (SVZ). Scale bar 5 *µ*m. (**c**) Hierarchical clustering of all single cells in (b) based on the Log2 transformation of mRNA transcript counts for these 7 genes. (**d**) Spatial illustration of molecular subgroups in mouse telencephalon neural progenitors identified in (c). Cluster color scheme is the same as it in (c).

## Discussion

Here, we present the HuluFISH as a new framework for smFISH. HuluFISH has the capability to enzymatically ligate multiple fluorophores to probes, which are designed based on their hybridization efficiency. And this new approach allows us to simultaneously detect genes with the multiplexity that increases exponentially with the number of available microscopy lasers and fluorophore types. With 4 to 5 color channels, it is possible to image 15 to 31 genes in one round of hybridization, which will fulfill a large number of experimental needs in detecting multiple RNA species, without resorting to multiple-step sequential hybridizations or super-resolution microscopy. HuluFISH labeling method is compatible with any other FISH related techniques. In particular, SeqFISH^4^ or MERFISH^5^ could employ HuluFISH labeling to either reduce the number of hybridization steps for fixed multiplexity or increase the multiplexity within their operational steps. Multiplexing *in situ* quantification of gene expression has become the next frontier in many fields for biomedical research. We believe the broad application of HuluFISH and its derivatives will greatly facilitate the discovery processes like cellular heterogeneity and precise gene expression regulation, in particular for the project like the Human Cell Atlas Initiative.

## Methods

### Cell culture and tissue section preparation

Mouse Hepa 1-6 cells were cultured in DMEM medium with 10% fetal bovine serum and 1 × penicillin/streptomycin. Hepa 1-6 cells were directly grown on coverslip without coating. Embryonic mouse brain tissue cryo-sections were cut at 6 to 10 µm from embryonic day 12.5 C57BL/6J mouse embryo embedded in Tissue-Tek O.C.T. (Sakura, 4583). Adherent Hepa 1-6 cells or cryo-sections were fixed with 4% formaldehyde in PBS for 10 min and then quenched with 135 mM glycine in PBS for 10 min at room temperature. Fixed cells were then washed once with PBS and permeabilized in 70% ethanol overnight at 4 °C. All water used for FISH related buffers was diethyl pyrocarbonate (DEPC) treated. After permeabilization cells were stored in cryoprotectant (25% glycerol, 25% ethylene glycol, 0.1M phosphate buffer, pH 7.4) at -20 °C until FISH staining.

### Probe design

smFISH probes based on the conventional design^3^ were implemented in an R script to select GSOs first with Primer3^11^ to get all possible GSOs without strong secondary structure from the input mRNA sequence using the standard condition for selecting the right_primer in Primer3. Then non-overlapping GSOs were selected with minimally 2 bp gap. For HuluFISH 1.0 probes, all GSOs from Primer3 were additionally calculated for their hybridization efficiency with DECIPHER package in R^12^ under the condition used for staining. And the GSOs were filtered to have hybridization efficiency above 0.9 (maximally 1) and then non-overlapping HuluFISH GSOs were selected as before. For HuluFISH 2.0 GSOs, additional tag sequence was added to their 3’ end after their selection. Adaptor, tag for GSO and Hulu sequences were randomly generated in an R script and controlled for strong secondary structure by UNAfold^15^. Passed sequences were blasted against a local mouse and human transcript database (ensemble release 87) for less than or equal to 15 bp exact match.

### HuluFISH probe labeling and purification

HuluFISH was initially an acronym for Helix-stabilized, unbiased and ligated uni/multi-color probe for FISH. In search of a multicolor object such as rainbow and confetti to name this technology in an imagery fashion, we got the inspiration from a famous Chinese cartoon, Hulu Brothers (húlú is calabash in Chinese), where each of the seven protagonists was transformed from a calabash with a distinct rainbow color, much like the base color in HuluFISH multiplexing. Besides, calabash fruits on a winding vine resemble fluorophores on a helical HuluFISH probe. Additionally, húlú bears the image of life in Chinese culture: it is a container for elixir and a symbol for reproduction, coinciding with the intended use of HuluFISH in medicine and life science.

FISH GSOs and adaptor oligonucleotides were synthesized from Sigma with the lowest quality for purification (desalting). For the individual gene, GSOs were pooled together to have 100 *µ*M total oligonucleotide concentration. Fluorescent Hulu oligonucleotides were purchased from Eurofins Genomics with various dyes, including Atto dyes, Alexa dyes or Cy dyes. For HuluFISH 1.0, ligation was performed in T4 DNA ligase buffer (NEB, B0202S), with 30 *µ*M adaptor for HuluFISH 1.0, 3 µM GSOs and Hulu oligonucleotide, 25% PEG8000, 30 U/*µ*L T4 DNA ligase (NEB, M0202M). Ligation reaction mix was then incubated in a thermocycler, with 12 cycles of 37 °C 10 seconds / 16 °C 5 minutes. For HuluFISH 2.0, ligation reaction mix was prepared as HuluFISH 1.0 with some modifications, such as 16.7 *µ*M of GSOs, adaptor for HuluFISH 2.0 and Hulu 2.0 oligonucleotides, 50 U/*µ*L T4 DNA ligase. Then the ligation mix was left in the dark at room temperature for 2 hours. The ligation product was concentrated with 9 volumes of butanol and centrifuged as pellet at 20,000 g, 15 minutes at 4 °C. colorful labeled oligonucleotide pellet was washed once with 100% ethanol and spin down to remove ethanol, then resolubilized in loading buffer (8M Urea, 1 × TBE (Carl Roth, A118.1), 0.01% bromophenol blue and xylene cyanol). With 5 minute denaturing at 90 °C, oligonucleotides were loaded onto 15% Urea-PAGE gel (8M Urea, 1 × TBE, 15% Rotiphorese Gel 30 (Carl Roth, 3029.2), 0.05% ammonium persulfate, 0.05% tetramethylethylenediamine) pre-run at 300 V for 30 minutes. Running condition was usually 300 V, 30 minutes, or until the bromophenol blue reached the end. Gel bands with fluorescent dye-oligonucleotide conjugates were excised under the ambient light. Gel pieces were homogenized manually by microtube pestle (Sigma, Z359947-100EA), and then extracted with 500 *µ*L 10 mM TE buffer (pH 8.5, 10 mM tris(hydroxymethyl)aminomethane (Tris), 1 mM Ethylenediaminetetraacetic acid (EDTA)) at room temperature overnight, protected from light by wrapping in aluminum foil. The extracted oligonucleotides in TE were concentrated again by butanol and washed once by ethanol like before. The final pellets were dried in dark at room temperature for 5-10 minutes, and then resolubilized in H_2_O. The concentration was determined by Nanodrop One (ThermoFisher) as ssDNA.

### FISH probe staining and imaging

HuluFISH probe mix was adjusted to 10 nM for each single oligonucleotide in hybridization buffer (2 × SSC (saline-sodium citrate), 10 % (w/v) dextran sulfate, 10% (v/v) formamide, 1 mg/mL tRNA (Roche, 10109541001), 2 mM ribonucleoside vanadyl complex (NEB, S1402S), 0.2 mg/mL BSA). Gapdh-Quasar570 probe was purchased from Biosearch Technology, resuspended and used for the staining as instructed from the manufacturer. Hybridization was performed in a water bath at 30 °C overnight, with the sample faced down on the parafilm. Cells on coverslip or tissue sections on glass slide were washed with washing buffer (2 × SSC, 10 % (v/v) formamide, 0.1 % (w/v) Tween-20) at 37 °C for 6 × 10 minutes. The last washing step included 0.5 *µ*g/mL DAPI (4’,6-diamidino-2-phenylindole) for nuclei staining. The sample was mounted in ProLong Gold Antifade (ThermoFisher, P10144), and cured overnight. The sample then was either imaged on a widefield microscope (Zeiss Cell Observer) with 200 ms, 950 ms and 5000 ms for 405nm, 488 nm and 561 nm channel, or on a confocal microscope with Airyscan (Zeiss LSM800, equipped with 405, 488, 561, and 640 nm laser) with maximal laser power (c.a. 5%) in each channel. The sample was scanned with Airyscan technology with the optimal settings provided by ZEN software.

### Image analysis

Except for the nuclear outline manually defined in ImageJ, all the image analysis was performed in R, and majorly based upon the package EBImage^16^. All intensity threshold values were based on the arbitrary units generated from Zeiss Airyscan and thus not specified in the following description. FISH dot identification relied on 2D local maxima identification and alignment. Initially for each frame, 2D maxima above a low threshold value were identified. Each 2D local maximum regarded its projection on the neighboring z-slices for alignment: those that fall within 0.08 μm were assigned to the same FISH dot. The pixels with maximal intensities (pseudo-3D-maxima) for identified FISH dots were extracted for further analysis.

Signal-to-noise ratio (SNR) and contrast were generated adaptively for each individual FISH dot. To this end, pixel values (local background) were taken from a square centered around the pseudo-3D-maxima, excluding all circular regions covering the PSF (point spread function) for 2D maxima on the same plane. Contrast is defined as the ratio of the maximal intensity and the mean of its local background values; SNR, as traditionally defined, equals to maximal intensity divided by the standard deviation of local background values.

For color decoding in samples with Hulu-probe for multiple genes, the presence of fluorophore on each channel was initially separately determined. Dual or triple color coding was assigned when FISH dots from different channels co-localized within 0.08 μm. The single color assignment required thresholding with a higher intensity, given there were three copies of fluorophores in the single-color Hulu-probe. Nuclei were manually segmented on the maximum intensity projected image in ImageJ. Without the assistance of membrane immunostaining, each identified FISH dot was assigned to its closest nuclei.

### Statistical analysis

Wilcoxon two samples test was used for evaluating the significance of our probe design based on HybEff and the conventional one based on Tm. The p-value is indicated in the corresponding figure legend. Single cell gene set expression data were hierarchically clustered using Euclidean distance and shown as a heatmap. 5 clusters were retrieved from 58 cells by cutting the dendrogram tree.

## Author Contributions

Y.S.C and H.K.L conceived and designed the project. Y.S.C performed most of the experiments. Y.Z helped with image acquisition and analysis, K.H, P.Z. G.B and H.A helped with FISH staining H.K.L supervised the project. Y.S.C, Y.Z, and H.K.L analyzed the results and wrote the manuscript.

## Acknowledgement

We thank Sven Poppelreuther at Carl Zeiss Imaging center in the DKFZ for providing critical suggestions in using Airyscan technology. We thank Bernd Fisher for generously providing R code for local maxima. This work was supported by the Helmholtz Association (VH-NG-702), the Deutsche Forschungsgemeinschaft (LI 2140/1-1), the Deutsche Krebshilfe (110226), the Helmholtz Alliance Preclinical Comprehensive Cancer Center (PCCC), the Chica Heinz Schaller foundation, the EMBO young investigator program, and the ERC (European Research Council) consolidator grant (647055) to H.K.L.

